# Development of SARS-CoV-2 mRNA vaccines encoding spike N-terminal and receptor binding domains

**DOI:** 10.1101/2022.10.07.511319

**Authors:** Guillaume B.E. Stewart-Jones, Sayda M. Elbashir, Kai Wu, Diana Lee, Isabella Renzi, Baoling Ying, Matthew Koch, Caralyn E. Sein, Angela Choi, Bradley Whitener, Dario Garcia-Dominguez, Carole Henry, Angela Woods, LingZhi Ma, Daniela Montes Berrueta, Laura E. Avena, Julian Quinones, Samantha Falcone, Chiaowen J. Hsiao, Suzanne M. Scheaffer, Larissa B. Thackray, Phil White, Michael S. Diamond, Darin K. Edwards, Andrea Carfi

## Abstract

With the success of mRNA vaccines against coronavirus disease 2019 (COVID-19), strategies can now focus on improving vaccine potency, breadth, and stability. We present the design and preclinical evaluation of domain-based mRNA vaccines encoding the wild-type spike-protein receptor-binding (RBD) and/or N-terminal domains (NTD). An NTD-RBD linked candidate vaccine, mRNA-1283, showed improved antigen expression, antibody responses, and stability at refrigerated temperatures (2-8°C) compared with the clinically available mRNA-1273, which encodes the full-length spike protein. In mice administered mRNA-1283 as a primary series, booster, or variant-specific booster, similar or greater immune responses and protection from viral challenge were observed against wild-type, beta, delta, or omicron (BA. 1) compared with mRNA-1273 immunized mice, especially at lower vaccine dosages. These results support clinical assessment of mRNA-1283 (NCT05137236).

**One Sentence Summary:** A domain-based mRNA vaccine, mRNA-1283, is immunogenic and protective against SARS-CoV-2 and emerging variants in mice.

---

In response to the coronavirus disease 2019 (COVID-19) pandemic, several vaccines were developed, including the mRNA-lipid nanoparticle (LNP) vaccines mRNA-1273 (Spikevax; Moderna, Inc., Cambridge, MA, USA) and BNT162b2 (Comirnaty; Pfizer Inc, New York, NY, USA; BioNTech Manufacturing GmbH, Mainz, Germany). These vaccines were efficacious in clinical trials, with continued real-world vaccine effectiveness (VE) against severe acute respiratory syndrome coronavirus 2 (SARS-CoV-2)–related infection, hospitalization, and death (*1–4*) and also against infection with SARS-CoV-2 variants beta (B.1.351), delta (B.1.617.2), and to a lesser extent, omicron (B.1.1.529)(*5*). The success of mRNA-based COVID-19 vaccines provides the opportunity to develop new vaccines focusing on dose-sparing, improved stability, and antigens tailored to key neutralization sites.

Most COVID-19 vaccines focus on the spike (S) protein trimer as the primary immunogen given its abundance on the virion surface and essential role in viral entry which makes it a target for neutralizing antibodies (nAbs) (*6*). The S protein is a type 1 fusion glycoprotein comprised of a C-terminal transmembrane region and an extracellular domain composed of the S1 and S2 subunits (**Fig. 1A-B**) (*7*). During priming for cellular entry, the S protein undergoes serine protease-mediated cleavage at the S1/S2 site, facilitating the binding of the receptor binding domain (RBD) to angiotensin-converting enzyme 2 (ACE2) and resulting in the successive fusion of viral and host membranes (*7–9*). mRNA-1273 encodes the full-length S protein containing 2 proline mutations within the S2 region (S-2P), resulting in a prefusion-stabilized S trimer, eliciting robust immunogenicity (*7, 10*).

**Figure 1.**
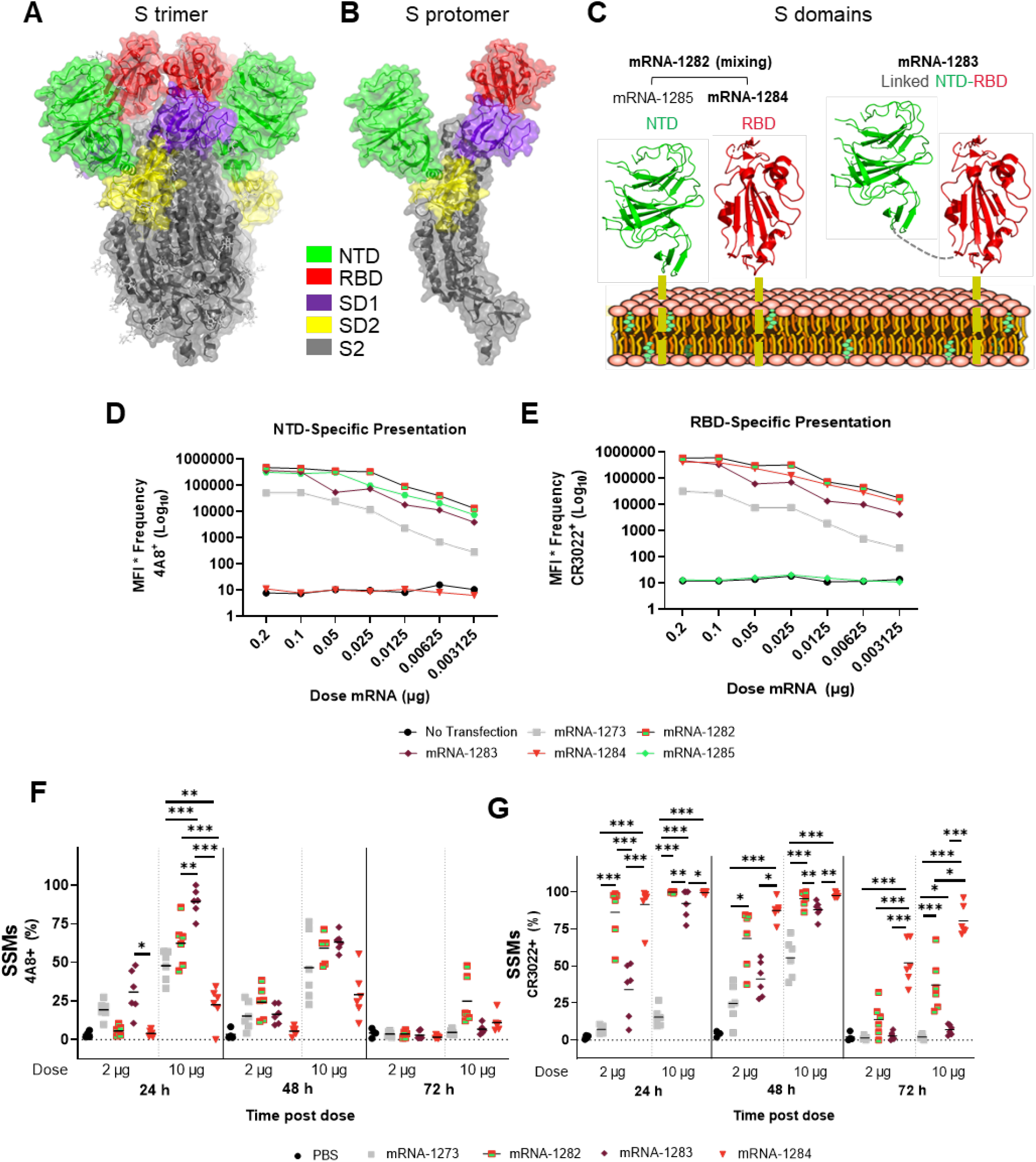
Structural design of novel mRNA vaccines and antigen presentation. Structural model of the trimeric S protein (**A**) and the S protein protomer (**B**) with domains highlighted. Vaccine domains with the transmembrane anchor (yellow dash) and peptide linkage (grey dash). (**C**) Average fold differences of MFI Freq for NTD (4A8; **D**) and RBD (CR3022; **E**) at 24 hours in HEK239T cells. Flow cytometry results for antigen expression in SSMs for RBD (CR3022; **F**) and NTD (4A8+; **G**) after 24, 48, and 72 hours; N=6; hypothesis testing was conducted using Bayesian information followed by the multivariate *t* distribution method. *, *P*<0.05, **, *P*<0.01, and, *** *P*<0.001.

Additional COVID-19 vaccine strategies have focused on including only the major antigenic domains (i.e., RBD and N-terminal domain [NTD]) to potentially increase protein expression and enhance immune responses to regions of the S protein that are targets of potent nAbs (*11–14*); the latter is especially important, as several variants including B.1.351, B.1.617.2, and B.1.1.529 contain multiple RBD and NTD mutations associated with immune evasion (*15, 16*). Additionally, it has been noted that immune imprinting by other coronavirus infections may impact SARS-CoV-2–specific responses because of shared S-protein domains across *Coronaviridae;* a SARS-CoV-2–specific domain vaccine could potentially overcome such issues (*17*). Furthermore, by only encoding for the major antigenic domains versus the full-length S protein, the reduced mRNA length could potentially improve vaccine shelf-life (*18*). Consequently, the adaptable nature of the mRNA vaccine platform is suited to the design and evaluation of domain-based vaccines that can be rapidly modified to encode variant-specific mutations.

Herein, we present the design and preclinical evaluation of multiple domain-encoded mRNA vaccines designed to express RBD and NTD (WA1/2020 prototype or variant-specific) as individual domains or in combination. We also evaluated the immunogenicity and protective activity of these vaccines against SARS-CoV-2 variant challenge in mice.

## Results

### Domain-based vaccine mRNA transfection demonstrates robust NTD and RBD protein expression in vitro

Four mRNA domain vaccines based on the WA1/2020 S sequence were developed encoding the RBD (mRNA-1284), NTD (mRNA-1285), RBD and NTD as 2 separate mRNAs mixed in a 1:1 ratio (mRNA-1282), and NTD-RBD connected by a G3SG3 linker (mRNA-1283; **Fig. 1C**). All mRNAs encoded a membrane anchor to facilitate cell surface expression and were encapsulated in LNPs.

HEK239T cells were transfected with mRNA-1273 or domain-vaccine mRNA, and surface expression of RBD and/or NTD was determined via flow cytometry. At 24 hours, a concentration-dependent increase in NTD and RBD expression was observed for all mRNAs (**Fig. 1D-E)**, reaching maximal expression at the highest dosage (0.2 μg). Cells transfected with domain vaccine mRNA consistently expressed higher levels of NTD or RBD than mRNA-1273-transfected cells at all dosages. A ≥ 14-fold increase in cell surface NTD and/or RBD expression at 24 hours post-transfection was observed for all mRNA administered at the lowest dose (0.003 μg) compared with mRNA-1273; similar observations were made for NTD and RBD expression at 48 and 120 hours (**Fig. S1**).

### Domain-based vaccine candidates demonstrate high levels of cell surface NTD and RBD expression in vivo

Based on the high NTD and RBD expression in cells by domain encoding mRNA, flow cytometry was used to determine domain expression in antigen presenting cells (APCs) of BALB/c mice administered mRNA-1273, mRNA-1282, mRNA-1283, or mRNA-1284 (all vaccines dosed at 2 μg or 10 μg); subscapular sinus macrophages (SSMs; CD19^-^, CD3^-^, IA/IE^+^, CD11b^+^, SIGN-R1^+^, CD169^+^), conventional dendritic cells (cDCs; CD19^-^, CD3^-^, Siglec H^-^, CD317^-^, CD11c^+^, IA/IE^+^), and plasmacytoid dendritic cells (pDCs; CD19^-^, CD3^-^, Siglec H^+^, CD317^+^) were isolated from draining inguinal lymph nodes for analysis of surface protein expression at 24, 48, and 72 hours post dose.

Cell surface NTD expression (**Fig. 1F**) on SSMs was highest at 24 hours in mice administered mRNA-1283 10 μg (mRNA-1273, *P*<0.001; mRNA-1284, *P*<0.001), with no significant differences observed at 48- and 72-hours compared with mRNA-1273. Low levels of surface NTD expression were observed on cDCs and pDCs (**Fig. S1D**).

For RBD, significant increases in cell surface expression (*P*<0.001) were observed in BALB/c SSMs 24 hours after administration of mRNA-1282, mRNA-1284, and mRNA-1283 compared with mRNA-1273 (10-μg doses; **Fig. 1G**); at 72 hours post-administration, significantly higher cell surface RBD expression was observed in the mRNA-1282 (*P*<0.001), mRNA-1283 (*P*<0.05), and mRNA-1284 (*P*<0.001) groups at 10-μg dose levels compared with mRNA-1273. Similar findings, although less pronounced, were observed for cDCs and pDCs (**Fig. S1C**).

### Domain-based vaccines are immunogenic in mice

We next evaluated the immunogenicity of single component domain vaccines in BALB/c mice by determining S-2P-, RBD- and NTD-specific immunoglobulin G (IgG) and nAb titers, as well as CD4^+^ and CD8^+^ T-cell cytokine responses to S1, S2, and RBD peptide pools. Mice (N=8-12/group) were administered either a 1-dose regimen (1 μg) of the single component vaccines mRNA-1273, mRNA-1283, mRNA-1284, or mRNA-1285 or a 2-dose regimen of the same vaccines (0.01, 0.1, or 1 μg) 21 days apart; immunogenicity was assessed at 21 days (pre-boost) and 36 days post dose 1 (15 days post-boost).

Binding and nAb titers were similar between mice administered a single 1-μg dose of the domain vaccines (mRNA-1283, mRNA-1284, and mRNA-1285) and mRNA-1273 and comparable to antibody responses observed in mice vaccinated with two 0.1 μg doses (**Fig. 2A and 2B**). All mRNA vaccines administered as a 2-dose regimen elicited similar S-2P specific IgG titers (**Fig. 2A**) in mice, with dose-dependent increases in responses observed. Highest nAb titers were generally observed in mice vaccinated with mRNA-1283 compared with the other vaccines (**Fig. 2B**). Domain-specific binding antibody titers were similar to those observed for S-2P specific responses for all vaccines; responses were consistently greater post-boost (Day 36; **Fig. S2B**) than pre-boost (Day 21; **Fig. S2A**). CD4^+^ and CD8^+^ T-cell responses were minimal at lower doses (0.1 μg) for all vaccines tested regardless of the peptide pool used (**Fig. S2C-D**). Compared with mRNA-1273–vaccinated mice, those vaccinated with mRNA-1284 and mRNA-1285 (both 1 μg) produced higher interferon gamma (IFN-γ) and tumor necrosis factor alpha (TNF-α) expression when CD4^+^ T-cells were stimulated with the S1 or RBD peptide pool; similar results were observed in CD8^+^ T-cells, when stimulated with the S1 or RBD peptide pool including those from mice vaccinated with mRNA-1283.

**Figure 2.**
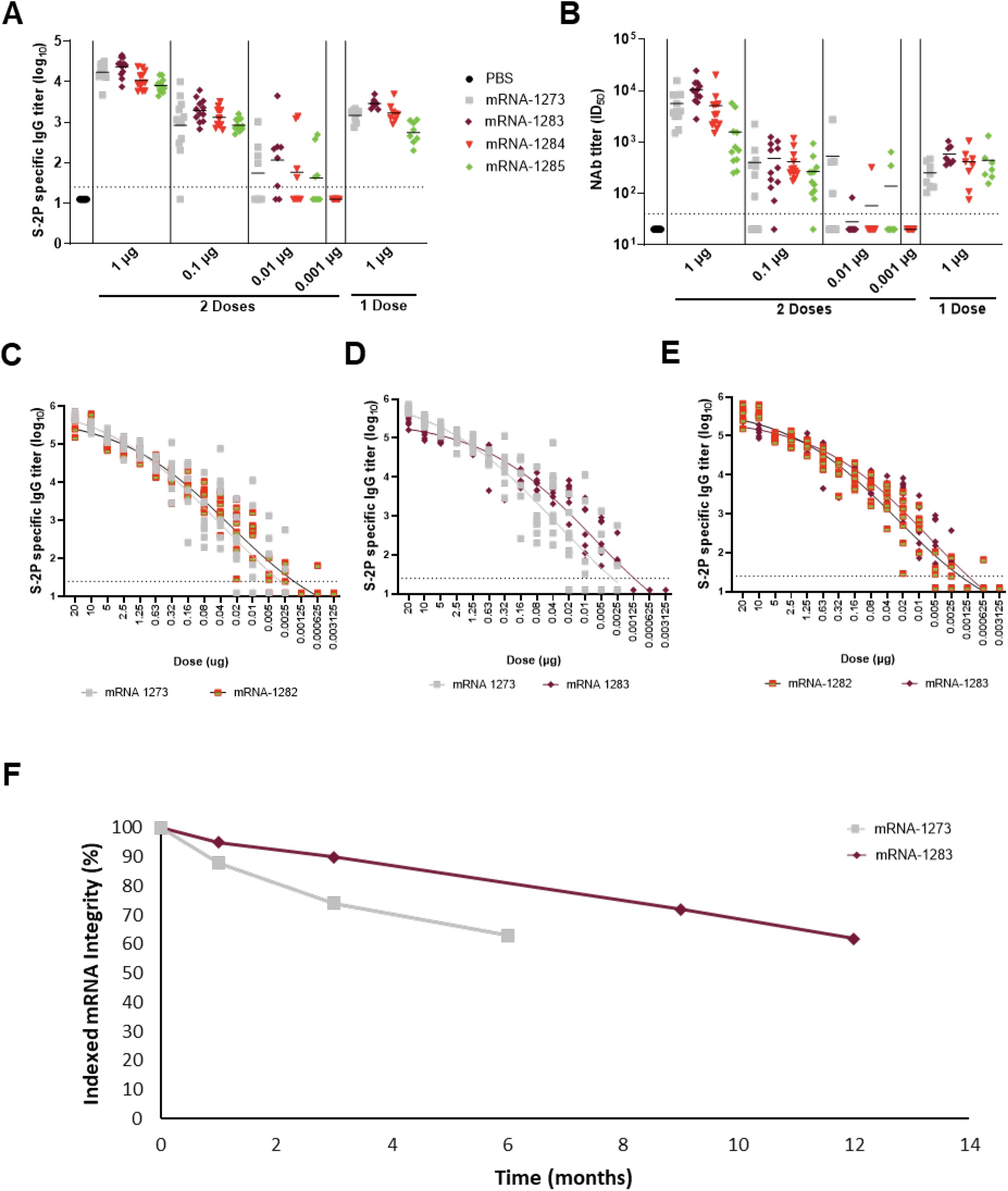
Immunogenicity, dose-sparing properties, and stability of domain-based mRNA vaccines. (**A-B**) S-2P-specific IgG titers and nAb titers for 1- or 2-dose domain vaccine regimens; N=8-12. (**C-E**) Dose response curves for RBD- and NTD-encoding vaccines. (**F**) Commercial representative GMP drug product lot used to evaluate mRNA degradation rates for mRNA-1273 and mRNA-1283 when stored at 2°-8°C. Values have been indexed to arbitrary units; N=8.

mRNA-1283 was further compared with a vaccine encoding the RBD and NTD as 2 separate sequences (mRNA-1282). Mice administered 2 doses of mRNA-1282 (**Fig. 2C**) or mRNA-1283 (**Fig. 2D**) had greater S-2P specific antibody responses at lower doses compared with mRNA-1273, despite the domain-based immunogens having significantly less surface area than the full-length mRNA-1273. However, when comparing the dose response curves of mRNA-1282 to mRNA-1283 (**Fig. 2E**), mRNA-1283 had greater S-2P specific antibody responses at lower doses. mRNA-1283 was selected for further evaluation, as antibody responses elicited by this vaccine trended higher than those observed for other vaccines evaluated, had comparable or greater T-cell responses than mRNA-1273, and encodes both the NTD and RBD.

### mRNA-1283 has improved refrigerated storage (2 -8°C) compared with mRNA-1273

Good manufacturing practice (GMP) stability studies were conducted using commercially representative clinical development lots of mRNA-1273 and mRNA-1283 (**Fig. 2F**) using reversed-phase ion-pair high-performance liquid chromatography to separate and quantitate full-length mRNA molecules from degradants. mRNA integrity of the shorter mRNA-1283 was compared to the full-sequence encoding mRNA-1273 at refrigerated temperatures over time. mRNA-1283 demonstrates improved 2-8°C storage stability, reaching 62% of its initial integrity at 12 months, compared with 63% integrity reached after only 6 months for mRNA-1273.

### mRNA-1283 primary series and variant-specific booster doses induce robust antibody responses in mice

We next evaluated the ability of mRNA-1283 to be adapted as a variant-specific vaccine against B.1.351. We previously showed that among multiple variants assessed prior to the emergence of B.1.1.529, nAb responses against B.1.351 had the greatest fold reduction (6.9-8.4) compared with D614G in mRNA-1273–vaccinated individuals (*19*). In response to B.1.351 emergence, mRNA-1273–based vaccines containing variant-specific S protein sequences or mRNA-1283–based vaccines containing variant-specific RBD/NTD mutations were evaluated in mice.

BALB/c mice (N=5/group) were first administered a 2-dose primary series (1 or 0.1 μg) of either mRNA-1273 or mRNA-1283 followed by a matched B.1.351-specific booster dose (mRNA-1273.351 for mRNA-1273–primed and mRNA-1283.351 for mRNA-1283–primed mice). D614G- and B.1.351-specific nAb titers were then determined (**Fig. 3A**; **Table S1**). In mRNA-1273–primed and mRNA-1283–primed mice, D614G-specific nAb titers decreased by 1.5-fold (**Fig. 3B**) and 1.9-fold (**Fig. 3C**), respectively, between Day 36 (14 days post dose 2) and pre-boost (Day 212); D614G-specific nAb titers were 2.9-fold and 2.3-fold higher in mRNA-1283 primed mice than the mRNA-1273 group on Days 36 and 212, respectively. In mice boosted with mRNA-1273.351 (1 μg), a 4.5-fold (D614G) and 15.3-fold (B.1.351) increase in nAb titers were observed 21 days post-boost (Day 233) compared with pre-boost (**Fig. 3D**); similar trends, albeit less pronounced, were observed for the 0.1-μg dose. In mice boosted with mRNA-1283.351 (1 μg), a 4.9-fold (D614G) and 40.6-fold (B.1.351) increase in nAb responses were observed post-boost compared with pre-boost. High fold increases in nAb titer against D614G (22.5-fold) and B.1.351 (41.2-fold) were observed in mice administered mRNA-1283.351 0.1 μg (**Fig. 3E**). Overall, a primary mRNA-1273 or mRNA-1283 series followed by a matched variant-specific booster increased D614G- and B.1.351-specific nAb titers compared with pre-boost in all groups evaluated. Fold differences in nAb titers between pre-boost and post-boost time points were greater for the mRNA-1283 groups than the mRNA-1273 groups, especially at lower doses (**Table S2**).

**Figure 3.**
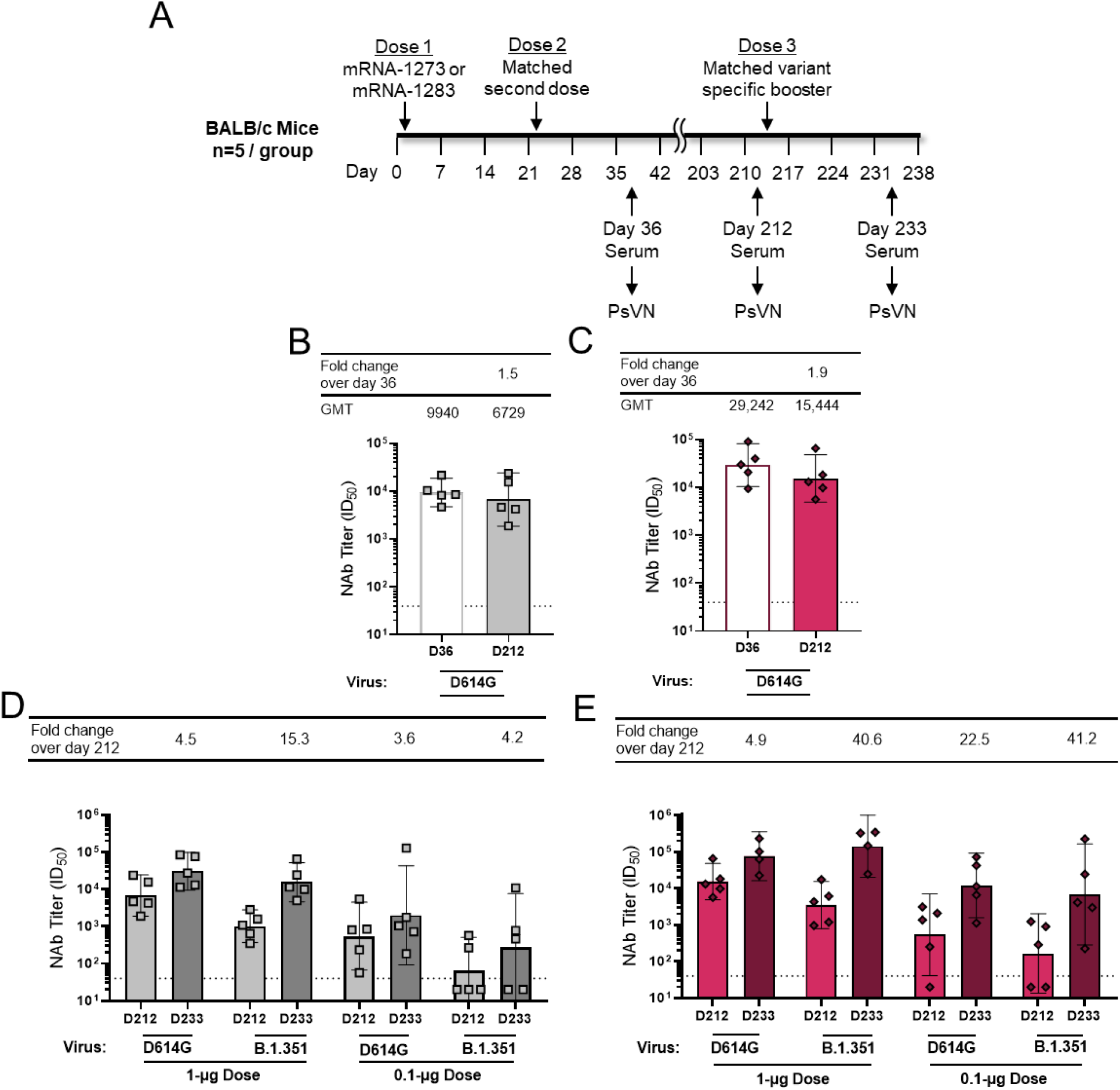
Immunogenicity in mRNA-1273– or mRNA-1283–primed mice receiving a matched B.1.351-specific booster. (**A**) Dosing schedule and sample collection. nAb titers in mice immunized with a 2-dose mRNA-1273 (1 μg; **B**) or mRNA-1283 (1 μg; **C**) primary series and against D614G and B.1.351 following mRNA-1273.351 (**D**) and mRNA-1283.351 (**E**) booster; N=5.

### mRNA-1283–based variant-specific vaccine boosters elicit greater nAb responses in mRNA-1273–primed mice than mRNA-1273–based boosters

Considering the widespread use of mRNA-1273 as a 2-dose primary series, it is necessary to evaluate how domain-based boosters compare with mRNA-1273–based boosters. Additionally, with the continuous emergence of SARS-CoV-2 variants, the adaptability of both vaccine types in inducing variant-specific immunogenic responses needs to be examined. Thus, nAb responses were assessed in mice that were administered a 2-dose primary mRNA-1273 series followed by a booster dose of mRNA-1273, mRNA-1283, or variant-specific mono-, stoichiometric di-, or trivalent vaccines in either mRNA-1273 or mRNA-1283 format (**Figure 4A and B**); all vaccines were administered at total mRNA 1.0 μg. Variant multivalent vaccines were administered due to their induction of nAb titers against multiple variants (*20*). Variant-specific vaccines included those specific to B.1.351 and B.1.617.2, as data published at study initiation reported lower vaccine-elicited nAb titers for these variants compared with D614G following primary mRNA-1273 vaccination (*19*). The S-protein mutations found in the different variants are illustrated in **Table S1.**

**Figure 4.**
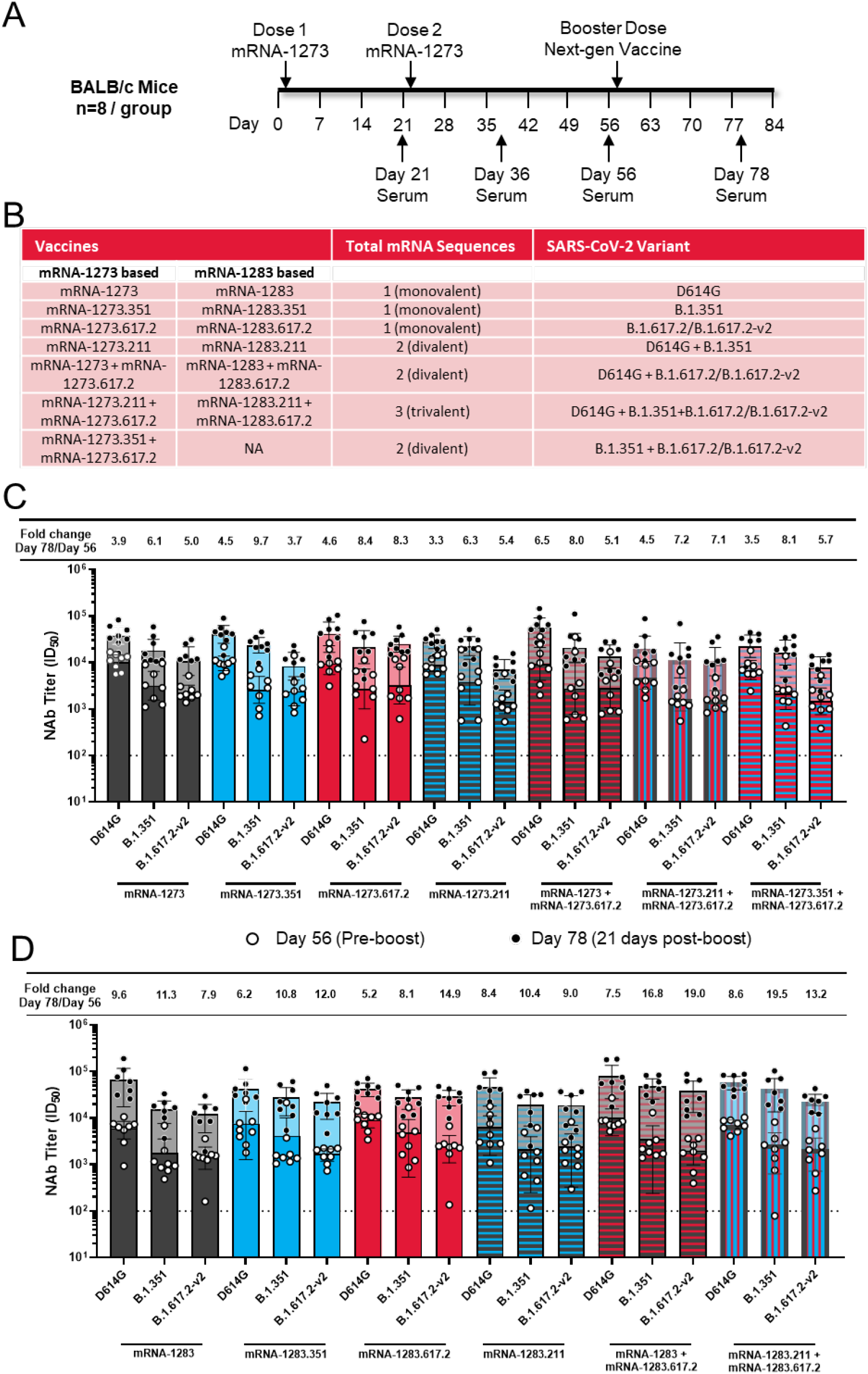
Immunogenicity in mRNA-1273–primed mice receiving an mRNA-1273 or mRNA-1283 variant-specific booster. (**A**) Dosing schedule and sample collection. (**B**) Details of vaccines investigated. (**C**) mRNA-1273 or (**D**) mRNA-1283 nAb titers for mice boosted with single- or multivalent vaccines; N=8.

At Day 78 (post-boost), up to 6.5-fold (D614G), 9.7-fold (B.1.351), and 8.3-fold (B.1.617.2) increases in nAb titers were observed in mice administered mRNA-1273–based boosters compared with Day 56 (pre-boost; **Fig. 4C**). The greatest fold increases in variant-specific nAb titers between pre- and post-boost was observed in mice administered mRNA-1273+mRNA-1273.617.2 against D614G (6.5-fold; divalent vaccine, **Fig. 4B**), mRNA-1273.351 against B.1.351 (9.7-fold), and mRNA-1273.617.2 against B.1.617.2-v2 (8.3-fold; **Fig. 4C** and **Table S3**). In mice administered an mRNA-1283–based booster, nAb titers increased up to 9.6-fold (D614G), 19.5-fold (B.1.351), and 19.0-fold (B.1.617.2) compared with pre-boost nAb titers (**Fig. 4D** and **Table S3**). The greatest fold increases in variant-specific nAb titers between pre- and post-boost was observed in mice administered mRNA-1283 against D614G (9.6-fold), mRNA-1283.211+mRNA-1283.617.2 against B.1.351 (19.5-fold; trivalent vaccine, **Fig. 4B**), and mRNA-1283+mRNA-1283.617.2 against B.1.617.2-v2 (19.0-fold; divalent vaccine; **Fig. 4B** and **4D**).

Overall, the greatest variant-specific nAb fold increases were observed in groups administered matched variant boosters for both mRNA-1273 and mRNA-1283, supporting the use of variant-matched boosters; fold changes in nAb titers between pre- and post-boost were more pronounced in mice vaccinated with mRNA-1283 than mRNA-1273.

### Mice vaccinated with mRNA-1283 were protected from B.1.1.529 (BA.1) challenge

SARS-CoV-2 variants with S protein mutations influence vaccine-induced immunity and protection from COVID-19 (*15, 16*). Accordingly, we assessed the ability of a primary mRNA-1283 series to confer protection against D614G (WA1/2020) or BA.1 in transgenic mice expressing human ACE2 (K18-hACE2), an animal model used to study severe COVID-19 (*21, 22*); for comparison purposes, figures include previously published time-matched control data from K18-hACE2 mice immunized with control mRNA and mRNA-1273, performed concurrently with the mRNA-1283 studies (*23*). Protection from BA. 1 was assessed as it is antigenically distant from the wild-type (D614G) and contains more than 30 S-protein mutations, many within the RBD and NTD (**Table S1**). K18-hACE2 mice were administered a primary series of either mRNA-1273 or mRNA-1283 (5 or 0.1 μg; 21 days apart), followed by challenge with 10^4^ focus-forming units (FFUs) of D614G or BA. 1 on Days 56-57; nAb titers were also evaluated using focus reduction neutralization tests with authentic SARS-CoV-2 viruses (**Fig. 5A**).

**Figure 5.**
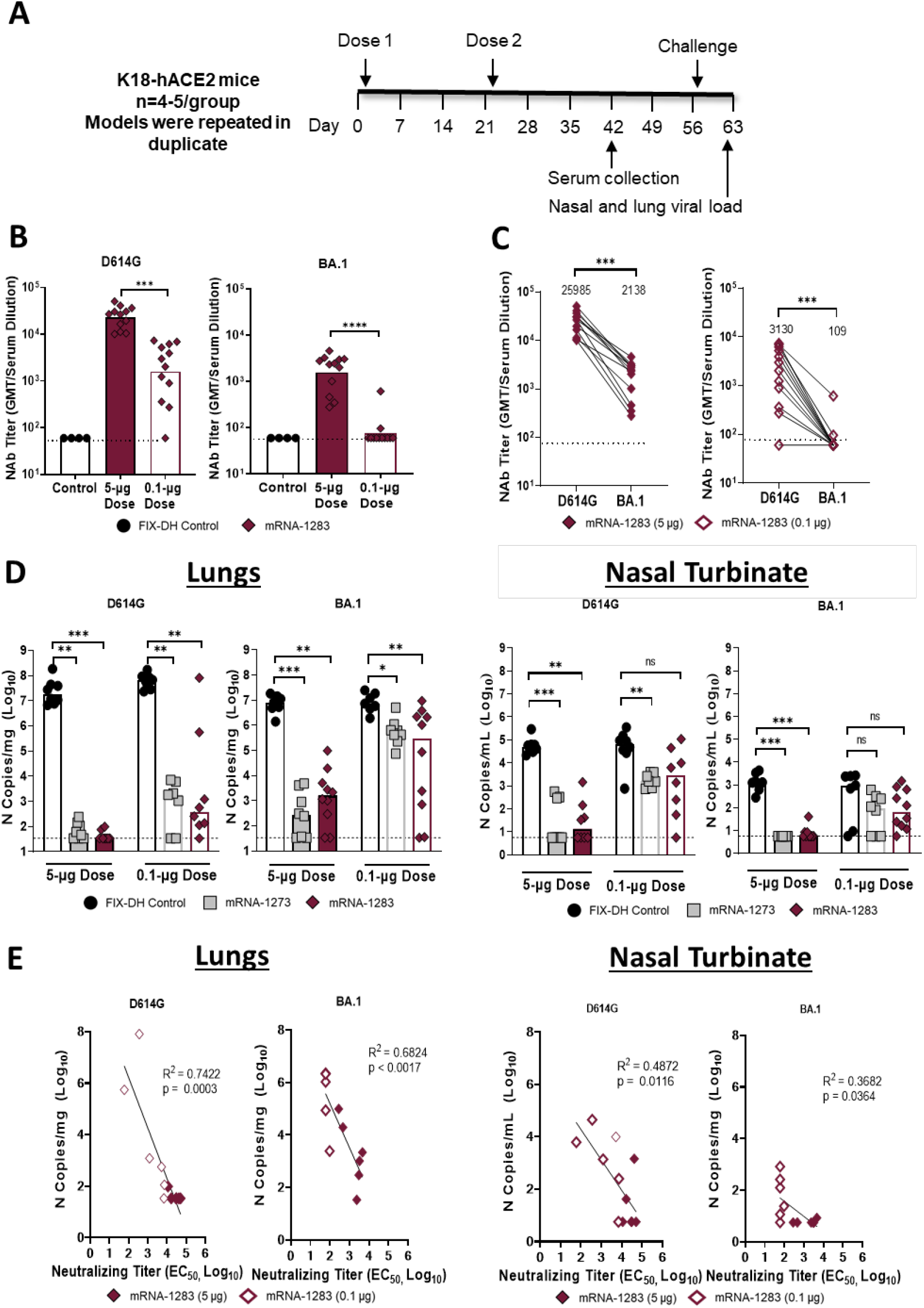
Immunogenicity and challenge with D614G and BA.1 in K18-hACE2 mice administered a primary series of mRNA-1273 or mRNA-1283. (**A**) Dosing schedule and sample collection. (**B**) nAb titers in mRNA-1283**–**vaccinated mice (N=8-10). (**C**) nAb titer for D614G and BA.1 in mRNA-1283**–**vaccinated mice (0.1 μg and 5 μg). GMTs are above each graph. (**D**) Lung or nasal viral *N* copies in mRNA-1273 or mRNA-1283**–**vaccinated mice (0.1- and 5-μg dose) challenged with D614G and BA.1 (N=8-10 per group; 2 repeat experiments). (**E**) Correlation between nAb titers and lung and nasal turbinate *N* copy numbers in mRNA-1283**–** vaccinated mice (0.1 μg and 5 μg); N=8-10 (N=4-5 per group; 2 repeat experiments). The data from control mRNA and mRNA-1273 groups are published but were performed concurrently with the mRNA-1283 studies and thus are provided in this figure for reference (*23*). Hypothesis testing was performed using the (**B, D**) Kruskal-Wallis ANOVA and Dunn’s test; (**C**) Wilcoxon matched-pairs signed rank test; (**E**) Pearson’s correlation and R^2^ value. For this figure, boxes illustrate geometric mean titer and dotted lines represent the limit of detection. ns, non-significant, *, *P*<0.05, **, *P*<0.01, and, ***, *P*<0.001.

Significantly greater nAb titers against BA. 1 (*P*<0.001) were observed in mice vaccinated with the 5-μg dose of mRNA-1283 versus mRNA-1283 0.1-μg dose (**Fig. 5B** and **Fig. S3**). nAb titers against BA. 1 were ~12-fold and ~29-fold lower at the 5-μg and 0.1-μg dose levels, respectively, compared with D614G (**Fig. 5C**).

After challenge with either WA1/2020 D614G or BA.1, viral burden was significantly lower in the lungs, nasal turbinate (**Fig. 5D**), and nasal washes (**Fig. S4**) of mice vaccinated with the 5 μg dose of mRNA-1283 compared with the control. D614G and BA. 1 viral *N* gene RNA copies were similarly low in mRNA-1273 and mRNA-1283–vaccinated mice. A lower level of protection was observed against BA.1 than D614G, regardless of vaccine or dose. When comparing nAb titers to viral *N* gene RNA copies in mice vaccinated with mRNA-1283, a strong inverse correlation was observed between nAb titers and viral *N* copies for both viruses (**Fig. 5E**); a similar, but less pronounced effect was observed in the nasal turbinate samples.

## Discussion

In this study, several SARS-CoV-2 domain-based mRNA vaccines demonstrated substantial *in vitro* and *in vivo* antigen expression and binding antibody responses, with mRNA-1283 (expressing NTD and RBD) selected for further evaluation. Specifically, among the domain based RNAs tested herein, a 2-dose primary series of mRNA-1283 in mice administered 21 days apart elicited higher antibody responses at lower doses than mRNA-1273. Additionally, variant-specific mRNA-1283 boosters produced greater increases in nAb titers versus mRNA-1273-based boosters against B.1.351 (≤19.5-fold vs ≤9.7-fold, respectively) and B.1.617.2 (≤19.0-fold vs ≤8.3-fold, respectively). Finally, K18-hACE2 transgenic mice administered a 2-dose mRNA-1283 series and challenged with D614G or BA. 1 5 weeks later were protected from both strains. Overall, these results, along with the potential of extended product shelf life, support the continued development of mRNA-1283 as a next generation SARS-CoV-2 vaccine.

Initial *in vitro* characterization revealed that domain-based vaccine mRNAs had greater antigen expression at all doses and time points evaluated compared with mRNA-1273. Robust vaccine-induced immunity is dependent on antigen presentation by APCs such as macrophages and DCs (*24, 25*). All 2-dose primary series vaccines tested had peak RBD and NTD cell surface expression in macrophages and DCs by 24 hours, with expression observed through 72 hours post-immunization, demonstrating the ability of these vaccines to induce rapid antigen expression lasting several days at levels greater than mRNA-1273, even at lower doses.

Immunogenicity assessments initially focused on mRNA-1284, mRNA-1285, and mRNA-1283 to ascertain NTD and RBD-specific immune responses generated within the context of vaccines encoding single mRNA sequences. These vaccines when administered as a 2-dose series in mice induced high levels of anti-S-2P IgG and nAb titers, with the highest nAb titers observed in the mRNA-1283 immunized group for doses >0.01 μg, consistent with published reports identifying NTD and RBD as principal antigenic targets following vaccination or natural infection (*14, 26*). Although the immunogenic surface area of domain-based vaccines is less than that of the full-length spike protein, similar or greater anti-S2P, -RBD, and -NTD IgG responses were observed for mRNA-1283 compared with mRNA-1273. Additionally, domain-based vaccines induced similar or greater T-cell cytokine responses compared with an mRNA-1273 primary series in mice, potentially due to greater surface antigen expression as was observed with mRNA-1284. These present important findings, as both humoral and cellular immunity mediate viral clearance, protect against variant-specific antibody escape, and confer durable protection against disease (*27–29*).

Further comparisons revealed that mRNA-1283 induced greater binding antibody responses in mice at lower doses compared with mRNA-1282 (RBD and NTD sequences in a 1:1 ratio) and mRNA-1273. These properties make mRNA-1283 an attractive candidate to advance for clinical evaluation as lower doses may limit reactogenicity as compared with mRNA-1273 without affecting immunogenicity and protection (*30, 31*). Additionally, mRNA integrity studies indicated that mRNA-1283 had a greater percentage of intact mRNA when stored at 5°C compared with mRNA-1273, which could assist with global COVID-19 vaccine storage and distribution strategies.

As new SARS-CoV-2 variants emerge, assessing the performance of vaccines against these strains is essential. Throughout this study, emerging variants of concern guided our experiments. Initial analyses focused on B.1.351, as this variant was previously shown to have substantially lower nAb titers compared with D614G in mRNA-1273–vaccinated individuals (*19*). Following a primary series of mRNA-1283, a booster dose of mRNA-1283.351 resulted in greater fold increases in D614G- and B.1.351-specific nAb titers compared with 2-dose mRNA-1273 followed by an mRNA-1273.351 booster; these findings were more pronounced at lower dose levels (0.1 μg versus 1 μg). Additionally, multiple mRNA-1283 variant-specific vaccines administered as a booster dose following a 2-dose primary mRNA-1273 series produced greater nAb titers than the equivalent mRNA-1273 variant-specific boosters. Finally, it is crucial to assess the ability of new and updated COVID-19 vaccines to protect against emerging variants. As expected, mice vaccinated with a 2-dose primary mRNA-1283 series had lower nAb titers against BA. 1 than D614G; nonetheless, post-challenge, significantly lower D614G and BA. 1 viral titers were observed compared with control groups. In general, nAb titers after immunization and viral copy levels post-challenge were comparable between mRNA-1283- and mRNA-1273–vaccinated mice. Taken together, these findings indicate that immune responses restricted to the RBD and NTD were sufficiently protective. Additionally, as these observations in mice remained true even at the lower dose tested (0.1 μg), they further support the dose-sparing properties of mRNA-1283 to elicit strong immunogenic responses with potentially lower reactogenicity (*30, 31*).

Our study has several limitations. As these are preclinical studies in a small number of mice, there may be differences in humans. Additionally, B.1.351, B.1.617.2, and BA.1 were the representative variants tested at the time of this study, necessitating further studies with emerging variants of concern. Moreover, mouse challenge studies only assessed protection after a primary series of mRNA-1283; a booster of mRNA-1283 in mRNA-1273 primed animals would be more reflective of real-world scenarios.

In the face of emerging SARS-CoV-2 variants, adapting vaccination strategies and using platforms that provide rapid and global distribution is essential. As such, mRNA-1283 shows considerable promise as a vaccine that can elicit robust immune responses at low doses and potentially can be stored considerably longer under refrigeration conditions. The ongoing evaluation of mRNA-1283 in a phase 2 clinical trial in healthy adults (NCT05137236) will provide important data regarding the safety and immunogenicity of this vaccine.

## Supporting information

Stewart-Jones 2022 bioRxiv_Supplemental Information

## Acknowledgements

***General*.** We thank Michael Brunner and Dr. Michael Whitt for kind support on recombinant VSV-based SARS-CoV-2 pseudovirus production. We would also like to thank the *in vivo* pharmacology and preclinical production teams at Moderna for performing the immunizations of BALB/c mice, and mRNA production and LNP formulation, respectively. Medical writing and editorial assistance, under the direction of the authors, was provided by Wynand van Losenoord, MSc, of MEDiSTRAVA in accordance with Good Publication Practice (GPP3) guidelines and funded by Moderna.

## Author contributions

GBESJ, LBT, MSD, DKE, and AC contributed to conceptualization; GBESJ, SME, KW, DL, IR, MK, CES, AC, BW, DGD, AW, LM, LEA, JQ, DKE, and AC contributed to data acquisition; GBESJ, SME, KW, DL, IR, MK, CES, AC, BW, DGD, CH, AW, LM, DMB, LEA, JQ, CJH, LBT, MSD, DKE, and AC contributed to the formal analysis; MSD and AC contributed to funding acquisition; GBESJ, SME, KW, DL, IR, BY, MK, CES, AC, BW, DGD, CH, AW, LM, DMB, LEA, JQ, SF, SMS, DKE, and AC contribute to the investigation; GBESJ, SME, KW, DL, IR, BY, MK, CES, BW, DGD, AW, LM, JQ, SMS, and DKE contributed to the methodology; GBESJ, LBT, DKE, and AC overseed project administration; GBESJ, DKE, and AC provided resources; GBESJ, SME, KW, LBT, DKE, and AC provided supervision; GBESJ, SME, KW, DL IR, CES, LM, DKE, and AC assisted with validation; GBESJ, SME, AC, DGD, CJH assisted with visualization; GBESJ, MSD, DKE, and AC assisted with writing the original draft development; all authors reviewed and edited manuscript development and approved the final draft.

## Data and Material Availability

The authors declare that the data supporting the findings of this study are available within this article and its Supplementary Information.

## Materials and Methods

### Next-generation mRNA vaccine design and synthesis

Three unique domain-based recombinant mRNA vaccine candidates were generated containing the RBD (mRNA-1284), NTD (mRNA-1285), and covalently linked NTD-RBD (mRNA-1283), all fused to an influenza hemagglutinin transmembrane anchor. An additional vaccine candidate was developed comprising a 1:1 mixture of mRNA-1285 and mRNA-1284 (mRNA-1282). Vaccines encoding variant-specific S protein mutations were generated for mRNA-1273 (S protein mutations) and mRNA-1283 (RBD and NTD mutations) in accordance with the mutations listed in **Table S1**. Each mRNA sequence was synthesized using codon-optimized nucleotides *in vitro* via an optimized T7 RNA polymerase-mediated transcription reaction with complete replacement of uridine by N1m-pseudouridine. Within the DNA template used, the antigen coding sequence was contained within the open reading frame, flanked by a 5’ and 3’ untranslated region sequences, with transcription terminated via an encoded polyA tail. Resulting mRNA products were capped with a 5’-Cap 1 structure using vaccinia capping enzyme (New England Biolabs, Ipswitch, MA) and Vaccinia 2’ O-methyltransferase (New England Biolabs). The mRNA was purified utilizing oligo-dT affinity chromatography and encapsulated in an LNP through a modified ethanol-drop nanoprecipitation process, as described previously (*32–34*). In brief, ionizable, structural, helper, and polyethylene glycol lipids were combined with prepared mRNA in acetate buffer (pH 5.0; at a 5:2 ratio of lipids:mRNA). The mixture was neutralized with Tris-Cl (pH 7.5), combined with sucrose as a cryoprotectant, sterile filtered, and stored frozen at –70°C until further use. The drug product was deemed acceptable for *in vivo* studies subsequent to analytical characterization, which included the determination of particle size and polydispersity, encapsulation, mRNA purity, double-stranded RNA content, osmolality, pH, endotoxin, and bioburden.

### In vitro expression

HEK239T cells (ATCC CRL-11268) were cultured in Dulbecco’s modified Eagle’s medium (DMEM) supplemented with 10% fetal bovine serum (FBS) and glutamine (2 mM) and maintained in a humidified incubator at 37°C and 5% CO2. Cells were transfected with mRNA-1273, mRNA-1282, mRNA-1283, mRNA-1284, and mRNA-1285 at various concentrations using a TransIT mRNA transfection kit (Mirus Bio, Madison, WI). At 24 hours, 48 hours, and 120 hours post-transfection, cells were collected and resuspended in fluorescence-activated cell sorting (FACS) buffer (1 x phosphate buffered saline [PBS] supplemented with 2% FBS). Cell surface expression of S protein was detected by staining resuspended cells with CR3022 (10 μg/mL; RBD specific antibody) or 4A8 (10 μg/mL; NTD-specific antibody) in FACS buffer for 30 minutes on ice; monoclonal antibodies were synthesized from the protein sequence deposited in the protein databank (PDB ID: 6W41 and 7C2L, respectively). Cells were then washed twice with FACS buffer and incubated with Alexa Fluor 647 goat anti-human IgG in FACS buffer for 30 minutes on ice. LIVE/DEAD Fixable Aqua Dead Cell Stain (Invitrogen [Waltham, MA], cat. L34966) was used to assess viability. Data was collected via an LSR Fortessa flow cytometer (Becton Dickinson) and analysis was performed using FlowJo software (version 10.7.1).

### In vivo expression

BALB/c mice were intramuscularly administered 10-μg or 2-μg doses of mRNA-1273, mRNA-1282, mRNA-1283, or mRNA-1284. Thereafter, inguinal lymph nodes were harvested 24-, 48-, or 72-hours post-dose. The lymph nodes were injected with a digestion solution consisting of 250 μg/mL of Liberase (Fisher Scientific [Pittsburgh, PA], cat. 50-100-3341) and 50 μg/mL of DNase (Fisher Scientific, cat. PI89836) in Roswell Park Memorial Institute 1640 medium (RPMI) supplemented with 2% FBS and 1% penicillin/streptomycin antibiotic, incubated for 15 minutes at 37°C, and quenched with RPMI + 10% FBS. The lymph nodes were subsequently crushed through 70 μm MACS SmartStrainers (Miltenyi Biotec [Bergisch Gladbach, Germany], cat. 130-110-916) to generate mononuclear single-cell suspensions. Cells were plated and washed with PBS prior to staining with LIVE/DEAD Fixable Aqua Dead Cell Stain (Invitrogen, cat. L34966) for 30 minutes at 4°C. Thereafter, cells were washed twice with Fc stain buffer (PBS supplemented with 3% FBS and 0.05% sodium azide), resuspended in Fc Block (clone 2.4G2, Becton, Dickinson, and Company [BD, Franklin Lakes, NJ], cat. 553142) and incubated for 5 minutes at 4°C. Staining was then performed at 4°C for 30 minutes with CR3022 or 4A8 and a surface stain cocktail containing the following antibodies: CD169 PerCP/Cy5.5 (BioLegend (San Diego, CA], cat. 142410), SIGN-R1 PE (R&D Systems [Minneapolis, MN], cat. FAB1836P), I-A/I-E Alexa Fluor 700 (eBioscience [San Diego, CA], cat. 56-5321-82), CD317 Brilliant Violet 605 (BioLegend, cat. 127025), Siglec-H Brilliant Violet 421 (BD, cat. 566581), CD3 FITC (BioLegend, cat. 100306), CD19 FITC (BioLegend, cat. 115506), CD11b APC/Cy7 (BioLegend, cat. 101226), CD11c PE/Cy7 (BD, cat. 558079). Cells were washed twice with Fc stain buffer and stained with Alexa Fluor 647 goat anti-human IgG (Southern Biotech [Birmingham, AL], cat. 2016-31) in FACS buffer for 30 minutes at 4°C followed by 2 wash steps with FC stain buffer and resuspended in Fluorofix (BioLegend, cat. 422101) for 30 minutes at 4°C. Finally, cells were washed with FC stain buffer, filtered through a 96-well plate 30μm filter (Pall [Port Washington], cat. 8027) and were resuspended in FC stain buffer prior to data collection via an LSR Fortessa flow cytometer (BD). Analysis was done using FlowJo software (version 10.7.1). Subscapular sinus macrophages were identified as CD19^-^, CD3^-^, IA/IE^+^, CD11b^+^, SIGN-R1^+^, and CD169^+^; conventional dendritic cells were identified as CD19^-^, CD3^-^, Siglec H^-^, CD317^-^, CD11c^+^, and IA/IE^+^; plasmacytoid dendritic cells were identified as CD19^-^, CD3^-^, Siglec H^+^, and CD317^+^.

### Animal models

Animal experiments were carried out in compliance with all pertinent US regulations and policies. The Moderna, Inc. animal *in vivo* pharmacology review committee reviewed and approved all animal experiments. For *in vivo* protein expression studies, female BALB/c mice (42-56 days old; Charles River Laboratories, Wilmington, MA) were immunized via intramuscular injections at 0 and 28 days. Vaccines were diluted in PBS and administered as 0.001-μg (mRNA-1284 only), 0.01-μg, 0.1-μg, or 1-μg doses. Mice were injected intramuscularly with 100 μL of a vaccine (50 μL into each hind leg), with a matching second dose administered at Day 28. Sera for antibody were collected 21 days after the first immunization and 14 days after the second immunization.

For the B.1.351 booster dose study in mice, mRNA-1273 and mRNA-1283 constructs were updated (mRNA-1273.351 and mRNA-1283.351, respectively) to incorporate the S protein mutations identified in B.1.351 (**Table S1**). BALB/c mice (N=8) were immunized with mRNA-1273 or mRNA-1283 1 or 0.1 μg on Days 1 and 22, and the level and durability of the antibody responses were evaluated over the course of 7 months. Sera collection occurred on day 212, and a third dose of mRNA-1273.351 or mRNA-1283.351 1 or 0.1 μg was administered on Day 213. nAb titers were evaluated on Day 233, 21 days post-boosting.

For the B.1.351 and B.1.617.2 booster study in mice, mRNA-1273 and mRNA-1283 parent constructs were updated to selectively incorporate mutations found on the B.1.351 and B.1.617.2 S protein (**Table S1**). BALB/c mice (N=8-10) received a 2-dose priming series (1 μg; Day 1 and Day 22) of mRNA-1273, followed by a mRNA-1273 (1 μg), mRNA-1283 (1 μg), variant-specific updated mRNA-1273 (1 μg), or variant-specific updated mRNA-1283 (1 μg) at Day 57. Mice were also administered combination vaccines including mRNA-1273.211 (2 mRNAs encoding the mutations of D614G and B.1.351 encapsulated within a single LNP). Serum was collected at Days 21, 36, 56, and 78 for variant-specific nAb titer evaluation.

For mouse D614G and BA.1 challenge studies, heterozygous K18-hACE2 C57BL/6J mice (strain: 2B6.Cg-Tg(K18-ACE2)2Prlmn/J, Cat # 34860) obtained from The Jackson Laboratory (Bar Harbor, ME) were housed in groups and fed standard chow diets. Mice were administered a primary series of either mRNA-1273 or mRNA-1283 21 days apart. At Day 42, nAb titers were evaluated followed by challenging of animals against D614G or BA.1 on Day 56/57. Studies with K18-hACE2 mice were carried out in accordance with the recommendations in the Guide for the Care and Use of Laboratory Animals of the National Institutes of Health. Studies were approved by the Institutional Animal Care and Use Committee at the Washington University School of Medicine (St. Louis, MO; assurance number A3381–01). Virus inoculations were performed under anesthesia that was induced and maintained with ketamine hydrochloride and xylazine, and all efforts were made to minimize animal suffering. Experiments were neither randomized nor blinded.

### mRNA integrity analysis via reversed-phase ion-pair chromatography (RP-IP)

Sample separation was performed on a 2.1 × 100 mm DNAPac RP column (4-μm particle size; Thermo Fisher Scientific), with a flow rate of 0.35 mL/minute and column temperature set at 65°C. Mobile phase A consisted of dibutylammonium acetate (50 mM; TCI America, Portland, OR) and triethylammonium acetate (100 mM; Sigma-Aldrich, Burlington, MA) and mobile phase B consisted of 50% acetonitrile (Sigma-Aldrich), dibutylammonium acetate (50 mM), and triethylammonium acetate (100 mM). Separation was accomplished using a step-gradient, with an initial 1.5-minute hold at 25% B, a 3.0-minute gradient from 25% to 50% B, a 14.5-minute gradient from 50% to 56% B, and a 0.5-minute gradient and hold at 100% B.

### Mouse SARS-CoV-2 challenge

Seven-week-old female K18-hACE2 C57BL/6 mice were immunized 3 weeks apart with of mRNA vaccines (5 or 0.1 μg) in of PBS 50 μl via intramuscular injection in the hind leg. Animals were bled 4 weeks after the second vaccine dose for immunogenicity analysis. Five weeks after completing the primary series immunization, mice were challenged with 10^4^ FFU of WA1/2020 D614G or BA.1 SARS-CoV-2 strains via the intranasal route. In all experiments, animals were euthanized 6 days after challenge. Of note, the data from control mRNA and mRNA-1273 vaccination were published previously (*23*) but used for reference, as these studies were performed concurrently with mRNA-1283 vaccination as part of a larger multi-arm study cohort series. Thus, these experiments are time-matched, not historical comparisons.

Tissues were weighed and homogenized with zirconia beads in a MagNA Lyser instrument (Roche Life Science, Basel, Switzerland) in DMEM medium 1 mL supplemented with 2% heat-inactivated FBS. Tissue homogenates were clarified by centrifugation at 10,000 rpm for 5 minutes and stored at -80°C. RNA was extracted using the MagMax mirVana Total RNA isolation kit (Thermo Fisher Scientific) on the Kingfisher Flex extraction robot (Thermo Fisher Scientific). RNA was reverse transcribed and amplified using the TaqMan RNA-to-CT 1-Step Kit (Thermo Fisher Scientific). Reverse transcription was carried out at 48°C for 15 minutes, followed by 2 minutes at 95°C. Amplification was accomplished over 50 cycles as follows: 95°C for 15 seconds and 60°C for 1 minute. Copies of SARS-CoV-2 *N* gene RNA in samples were determined using a published assay (*35*).

### Enzyme-linked immunosorbent assay

Microtiter plates (96-well; Thermo Fisher Scientific) were coated with S-2P protein 1 μg/mL (Genscript, Piscataway, NJ), RBD protein 1 μg/mL (Sino Biological, Beijing, China), or NTD protein 2 μg/mL (Moderna, Inc.). After overnight incubation at 4 °C, plates were washed 4 times with PBS/0.05% Tween-20 and blocked using Superblock (Thermo Fisher Scientific) for 1.5 hours at 37°C. After washing, 5-fold serial dilutions of mouse serum were added (assay diluent-PBS + 0.05% Tween-20 + 5% goat serum). Plates were incubated for 2 hours at 37°C, washed and horseradish peroxidase (HRP)-conjugated goat anti-mouse IgG (Southern Biotech) was added at a 1:20,000 dilution (S-2P), or a 1:10,000 dilution (RBD, NTD) in assay diluent. Plates were incubated for 1 hour at 37°C and washed; bound antibody was detected with TMB substrate (SeraCare, Milford, MA). After incubation for 10 minutes at room temperature, the reaction was stopped by adding TMB stop solution (SeraCare) and the absorbance was measured at 450 nm. Titers were determined using a 4-parameter logistic curve fit in GraphPad Prism (Version 9; GraphPad Software, Inc., San Diego, CA) and was defined as the reciprocal dilution at approximate optical density (OD)450 = 1.0 (normalized to a mouse standard on each plate).

### Focus reduction neutralization test

Serial dilutions of sera were incubated with 10^2^ FFU of WA1/2020 D614G or BA.1 for 1 hour at 37°C. Antibody-virus complexes were added to Vero-TMPRSS2 cell monolayers in 96-well plates and incubated at 37°C for 1 hour. Subsequently, cells were overlaid with 1% (w/v) methylcellulose in minimal essential media. Plates were harvested 30 hours (WA1/2020 D614G) or 72 hours (BA.1) later by removing overlays and fixed with 4% paraformaldehyde in PBS for 20 minutes at room temperature. Plates were washed and sequentially incubated with an oligoclonal pool (SARS2-02, -08, -09, -10, -11, -13, -14, -17, -20, -26, -27, -28, -31, -38, -41, -42, -44, -49, -57, -62, -64, -65, -67, and -71 (*23*) of anti-S murine antibodies (including cross-reactive mAbs to SARS-CoV) and HRP-conjugated goat anti-mouse IgG (Sigma Cat # A8924, RRID: AB_258426) in PBS supplemented with 0.1% saponin and 0.1% bovine serum albumin. SARS-CoV-2–infected cell foci were visualized using TrueBlue peroxidase substrate (KPL, Noida, India) and quantitated on an ImmunoSpot microanalyzer (Cellular Technologies, Shaker Heights, OH).

### Recombinant vesicular stomatitis virus–based pseudovirus neutralization assay

Codon-optimized full-length S proteins of the original Wuhan-Hu-1 variant with D614G mutation (D614G) or B.1.351variant were cloned as described previously (*36*). Human lung carcinoma cells expressing human angiotensin-converting enzyme 2 (A549-hACE2-TMPRSS2 cells) were infected by pseudoviruses for 18 hours at 37°C. Assay plates were equilibrated to room temperature, and equal volume of One-Glo reagent (Promega, Fitchburg, WI) was added to culture medium for readout using a BMG PHERastar-FS plate reader. Serial dilutions of mouse sera (4-fold, 8 dilutions) were mixed with pseudovirus strains, which were previously diluted to 100,000 relative luciferase units (RLUs). Sigmoidal curves, taking average of triplicates at each dilution, were generated from RLU readings; 50% neutralization (IC50) titers were calculated considering uninfected cells as 100% neutralization and cells transduced with only virus as 0% neutralization.

### T-cell analysis

Whole spleens were collected from BALB/c mice on Day 36. A gentleMACS tissue dissociator (Miltenyi Biotec) was used to generate mononuclear single-cell suspensions from whole mouse spleens. Following tissue dissociation, the cells were sieved through a 70-μm filter. Cells from each mouse were resuspended in R10 media (Roswell Park Memorial Institute 1640 medium supplemented with L-glutamine, penicillin/streptomycin antibiotic, and 10% heat-inactivated FBS) and incubated at 37°C for 6 hours with protein transport inhibitors GolgiStop and GolgiPlug (BD Biosciences) and S glycoprotein peptide pools 1 μg/mL (JPT product PM WCPV-S-1) or RBD peptide pool 1 μg/mL (JPT product PM-WCPV-S-RBD). Control cells were incubated with no peptide stimulation. Cells were then washed with PBS prior to staining with LIVE/DEAD aqua fixable stain (Invitrogen) for 20 minutes at room temperature. Cells were subsequently washed with Fc stain buffer (PBS supplemented with 3% HI-FBS and 0.05% sodium azide) and resuspended in BD Fc Block (clone 2.4G2) for 5 minutes at room temperature. Staining was then performed at 4°C for 30 minutes with a surface stain cocktail containing the following antibodies: CD4 APC (BioLegend, cat. 100412, clone GK1.5), CD8 Alexa Fluor 700 (BioLegend, cat. 126618, clone YTS156.7.7), and CD44 BV421 (BD, cat. 563970, clone IM7). Cells were subsequently washed with Fc buffer, then fixed and permeabilized by using Cytofix/Cytoperm (BD Biocsiences, cat. 554714). After cells were washed, intracellular cytokine staining was performed at 4°C for 30 minutes using a cocktail of antibodies resuspended in 1x Perm/Wash buffer (BD Biosciences); intracellular antibodies included IFNγ APC-Cy7 (BioLegend, cat. 505850, clone XMG1.2), TNFα PE-Dazzle594 (BioLegend, cat. 506345, clone MP6-XT22), IL-2 BV711 (BioLegend, cat. 503837, clone JES6-5H4), IL-4 PE-Cy7 (BioLegend, cat. 504117, clone 11B11), IL-5 PE (BioLegend, cat. 504303, clone TRFK5), IL-9 PerCP-Cy5.5 (BioLegend, cat. 514112, clone RM9A4), IL-10 BV605 (BioLegend, cat. 505031, clone JES5-16E3), and IL-13 Alexa Fluor 488 (ThermoFisher, cat. 53-7133-82, clone eBio13A). Finally, cells were washed, filtered through a 96-well plate 30-μm filter (Pall), and were resuspended in Fc stain buffer prior to performing flow cytometry analysis on a LSR Fortessa flow cytometer (BD Biosciences) and analyzed using FlowJo software (version 10.7.1). Background cytokine expression in the no peptide condition was subtracted from that measured in the peptide pools for each individual mouse.

### Statistical modelling and hypothesis testing

To compare cell surface antigen expression *in vivo*, a linear mixed effect model was used to jointly model the percentage of cells expressing NTD and RBD by the various subdomain vaccine constructs. Included in the model are random intercepts accounting for animal-specific effect across cell types and surface antigens, in addition to covariates of hours post dosing, cell type, sell surface antigens, dose level, vaccine constructs, together with higher-order interactions of these covariates. R version 4.1.2 (*37*) was used for statistical modeling. Multiple comparisons were conducted using the emmeans package in R (*38*), with multivariate t adjustment at an alpha level of 0.05, except when noted otherwise.

## References

1. L. R. Baden et al., Efficacy and Safety of the mRNA-1273 SARS-CoV-2 Vaccine. N Engl J Med 384, 403–416 (2021).

2. F. P. Polack et al., Safety and Efficacy of the BNT162b2 mRNA Covid-19 Vaccine. N Engl J Med 383, 2603–2615 (2020).

3. K. Bruxvoort et al., Real-World Effectiveness of the mRNA-1273 Vaccine Against COVID-19: Interim Results from a Prospective Observational Cohort Study. Preprints with THE LANCET, (2021).

4. G. Chodick et al., The effectiveness of the TWO-DOSE BNT162b2 vaccine: analysis of real-world data. Clin Infect Dis, (2021).

5. B. Zeng, L. Gao, Q. Zhou, K. Yu, F. Sun, Effectiveness of COVID-19 vaccines against SARS-CoV-2 variants of concern: a systematic review and meta-analysis. BMC Med 20, 200 (2022).

6. L. Dai, G. F. Gao, Viral targets for vaccines against COVID-19. Nat Rev Immunol 21, 73–82 (2021).

7. D. Wrapp et al., Cryo-EM structure of the 2019-nCoV spike in the prefusion conformation. Science 367, 1260–1263 (2020).

8. M. Hoffmann et al., SARS-CoV-2 Cell Entry Depends on ACE2 and TMPRSS2 and Is Blocked by a Clinically Proven Protease Inhibitor. Cell 181, 271–280 e278 (2020).

9. D. J. Benton et al., Receptor binding and priming of the spike protein of SARS-CoV-2 for membrane fusion. Nature 588, 327–330 (2020).

10. K. S. Corbett et al., SARS-CoV-2 mRNA vaccine design enabled by prototype pathogen preparedness. Nature 586, 567–571 (2020).

11. J. Yang et al., A vaccine targeting the RBD of the S protein of SARS-CoV-2 induces protective immunity. Nature 586, 572–577 (2020).

12. N. Wang, J. Shang, S. Jiang, L. Du, Subunit Vaccines Against Emerging Pathogenic Human Coronaviruses. Front Microbiol 11, 298 (2020).

13. P. J. M. Brouwer et al., Potent neutralizing antibodies from COVID-19 patients define multiple targets of vulnerability. Science 369, 643–650 (2020).

14. G. Cerutti et al., Potent SARS-CoV-2 neutralizing antibodies directed against spike N-terminal domain target a single supersite. Cell Host Microbe 29, 819–833 e817 (2021).

15. W. T. Harvey et al., SARS-CoV-2 variants, spike mutations and immune escape. Nat Rev Microbiol 19, 409–424 (2021).

16. X. Zhang et al., SARS-CoV-2 Omicron strain exhibits potent capabilities for immune evasion and viral entrance. Signal Transduct Target Ther 6, 430 (2021).

17. T. Aydillo et al., Immunological imprinting of the antibody response in COVID-19 patients. Nat Commun 12, 3781 (2021).

18. T. C. Santiago, I. J. Purvis, A. J. Bettany, A. J. Brown, The relationship between mRNA stability and length in Saccharomyces cerevisiae. Nucleic Acids Res 14, 8347–8360 (1986).

19. A. Choi et al., Serum Neutralizing Activity of mRNA-1273 Against SARS-CoV-2 Variants. J Virol, JVI0131321 (2021).

20. K. Ghazvini, M. Keikha, Multivalent vaccines against new SARS-CoV-2 hybrid variants. Vacunas, (2022).

21. P. B. McCray, Jr. et al., Lethal infection of K18-hACE2 mice infected with severe acute respiratory syndrome coronavirus. J Virol 81, 813–821 (2007).

22. E. S. Winkler et al., SARS-CoV-2 infection of human ACE2-transgenic mice causes severe lung inflammation and impaired function. Nat Immunol 21, 1327–1335 (2020).

23. B. Ying et al., Boosting with variant-matched or historical mRNA vaccines protects against Omicron infection in mice. Cell 185, 1572–1587 e1511 (2022).

24. G. T. Rijkers et al., Antigen Presentation of mRNA-Based and Virus-Vectored SARS-CoV-2 Vaccines. Vaccines (Basel) 9, (2021).

25. I. A. Cockburn et al., Prolonged antigen presentation is required for optimal CD8+ T cell responses against malaria liver stage parasites. PLoS Pathog 6, e1000877 (2010).

26. C. O. Barnes et al., SARS-CoV-2 neutralizing antibody structures inform therapeutic strategies. Nature 588, 682–687 (2020).

27. R. Keeton et al., T cell responses to SARS-CoV-2 spike cross-recognize Omicron. Nature, (2022).

28. A. Kusnadi et al., Severely ill COVID-19 patients display impaired exhaustion features in SARS-CoV-2-reactive CD8(+) T cells. Sci Immunol 6, (2021).

29. A. Chandrashekar et al., Vaccine protection against the SARS-CoV-2 Omicron variant in macaques. Cell 185, 1549–1555 e1511 (2022).

30. M. N. Ramasamy et al., Safety and immunogenicity of ChAdOx1 nCoV-19 vaccine administered in a prime-boost regimen in young and old adults (COV002): a single-blind, randomised, controlled, phase 2/3 trial. Lancet 396, 1979–1993 (2021).

31. J. Mateus et al., Low-dose mRNA-1273 COVID-19 vaccine generates durable memory enhanced by cross-reactive T cells. Science 374, eabj9853 (2021).

32. K. J. Hassett et al., Optimization of Lipid Nanoparticles for Intramuscular Administration of mRNA Vaccines. Mol Ther Nucleic Acids 15, 1–11 (2019).

33. K. S. Corbett et al., Evaluation of the mRNA-1273 Vaccine against SARS-CoV-2 in Nonhuman Primates. N Engl J Med 383, 1544–1555 (2020).

34. J. Nelson et al., Impact of mRNA chemistry and manufacturing process on innate immune activation. Sci Adv 6, eaaz6893 (2020).

35. J. B. Case, A. L. Bailey, A. S. Kim, R. E. Chen, M. S. Diamond, Growth, detection, quantification, and inactivation of SARS-CoV-2. Virology 548, 39–48 (2020).

36. J. Pallesen et al., Immunogenicity and structures of a rationally designed prefusion MERS-CoV spike antigen. Proc Natl Acad Sci U S A 114, E7348–E7357 (2017).

37. R Core Team. (2021).

38. R. V. Lenth, emmeans: Estimated Marginal Means, aka Least-Squares Means. R package version 1.7.2. https://CRAN.R-project.org/package=emmeans. (2022).

