## Supplementary material for "Development of SARS-CoV-2 mRNA vaccines encoding spike N-terminal and receptor binding domains": Stewart-Jones 2022 bioRxiv_Supplemental Information

Stewart-Jones, et al.

**Table S1.** Variant-specific S protein mutations

| **Variant Name** | **Amino Acid Changes in Spike** |
|---|---|
| D614G | D614G |
| B.1.351 | L18F-D80A-D215G-ΔLAL242-244-R246I-K417N-E484K-N501Y-D614G-A701V |
| B.1.617.2 | T19R-G142D-E156G-F157del-R158del-L452R-T478K-D614G-P681R-D950N |
| B.1.617.2-v2 | T19R-T95I-G142D-E156G-F157del-R158del-L452R-T478K-D614G-P681R-D950N |
| BA.1 | A67V-Δ69-70-T95I-G142D/Δ143-145-Δ211-L212I-ins214EPE-G339D-S371L-S373P-S375F-K417N-N440K-G446S-S477N-T478K-E484A-Q493R-G496S-Q498R-N501Y-Y505H-T547K-D614G-H655Y-N679K-P681H-N764K-D796Y-N856K-Q954H-N969K-L981F |

**Table S2.** Neutralizing antibody titers (GMT) in mice administered a 2-dose primary series of mRNA-1273 or mRNA-1283 at different dose levels boosted with a matched B.1.351-specific vaccine.

| | **Neutralizing Antibody Titers (GMT)** | | | |
|---|---|---|---|---|
| | D614G | | B.1.351 | |
| **Primary Vaccine Series** | Day 212 | Day 233 | Day 212 | Day 233 |
| mRNA-1273 (0.1 µg) | 548 | 1985 | 66 | 274 |
| mRNA-1273 (1.0 µg) | 6729 | 30,386 | 1015 | 15,524 |
| mRNA-1283 (0.1 µg) | 538 | 12,091 | 167 | 6884 |
| mRNA-1283 (1.0 µg) | 15,444 | 76,399 | 3513 | 142,683 |

**Table S3.** Neutralizing antibody titers (GMT) in mice boosted with variant-specific mRNA-1273 and mRNA-1283 doses.

| Vaccine (Dose 3) | D614G | | B.1.351 | | B.1.617.2 | |
|---|---|---|---|---|---|---|
| | Day 56 (GMT) | Day 78 (GMT) | Day 56 (GMT) | Day 78 (GMT) | Day 56 (GMT) | Day 78 (GMT) |
| **mRNA-1273 parent sequence** | | | | | | |
| **mRNA-1273** | 10,310 | 40,483 | 3191 | 19,436 | 2308 | 11,457 |
| **mRNA-1273.351** | 9647 | 43,177 | 2579 | 25,136 | 2426 | 8865 |
| **mRNA-1273.617.2** | 9587 | 44,423 | 2716 | 22,852 | 3197 | 26,651 |
| **mRNA-1273.211** | 8933 | 29,693 | 3786 | 24,017 | 1422 | 7647 |
| **mRNA-1273 + mRNA-1273.617.2** | 9339 | 60,668 | 2761 | 22,226 | 2819 | 14,377 |
| **mRNA-1273.211 + mRNA-1273.617.2** | 4726 | 21,166 | 1650 | 11,865 | 1581 | 11,227 |
| **mRNA-1273.351 + mRNA-1273.617.2** | 6891 | 24,021 | 2165 | 17,490 | 1466 | 8379 |
| **mRNA-1283 parent sequence** | | | | | | |
| **mRNA-1283** | 4980 | 47,741 | 1217 | 13,738 | 1250 | 9900 |
| **mRNA-1283.351** | 5532 | 34,304 | 2162 | 23,250 | 1561 | 18,792 |
| **mRNA-1283.617.2** | 8296 | 42,747 | 3203 | 26,074 | 1872 | 27,973 |
| **mRNA-1283.211** | 4516 | 37,909 | 1157 | 11,996 | 1643 | 14,779 |
| **mRNA-1283 + mRNA-1283.617.2** | 8748 | 65,446 | 2585 | 43,371 | 1606 | 30,528 |
| **mRNA-1283.211 + mRNA-1283.617.2** | 6941 | 59,558 | 1588 | 30,895 | 1575 | 20,865 |

**Figure S1. Analysis of antigen expression.** Average fold difference (over dilution range) in MFI*Freq of NTD (mAb 118; **A**) and RBD (CR3022; **B**) at 48 hours (top) and 120 hours (bottom) in HEK239 T cells. FACS results for antigen expression in cDC and pDC for NTD (4A8+; **C**) and RBD (CR3022; **D**) after 24, 48, and 72 hours; N=6. *, $P<0.05$; **, $P<0.01$; and ***, $P<0.001$.

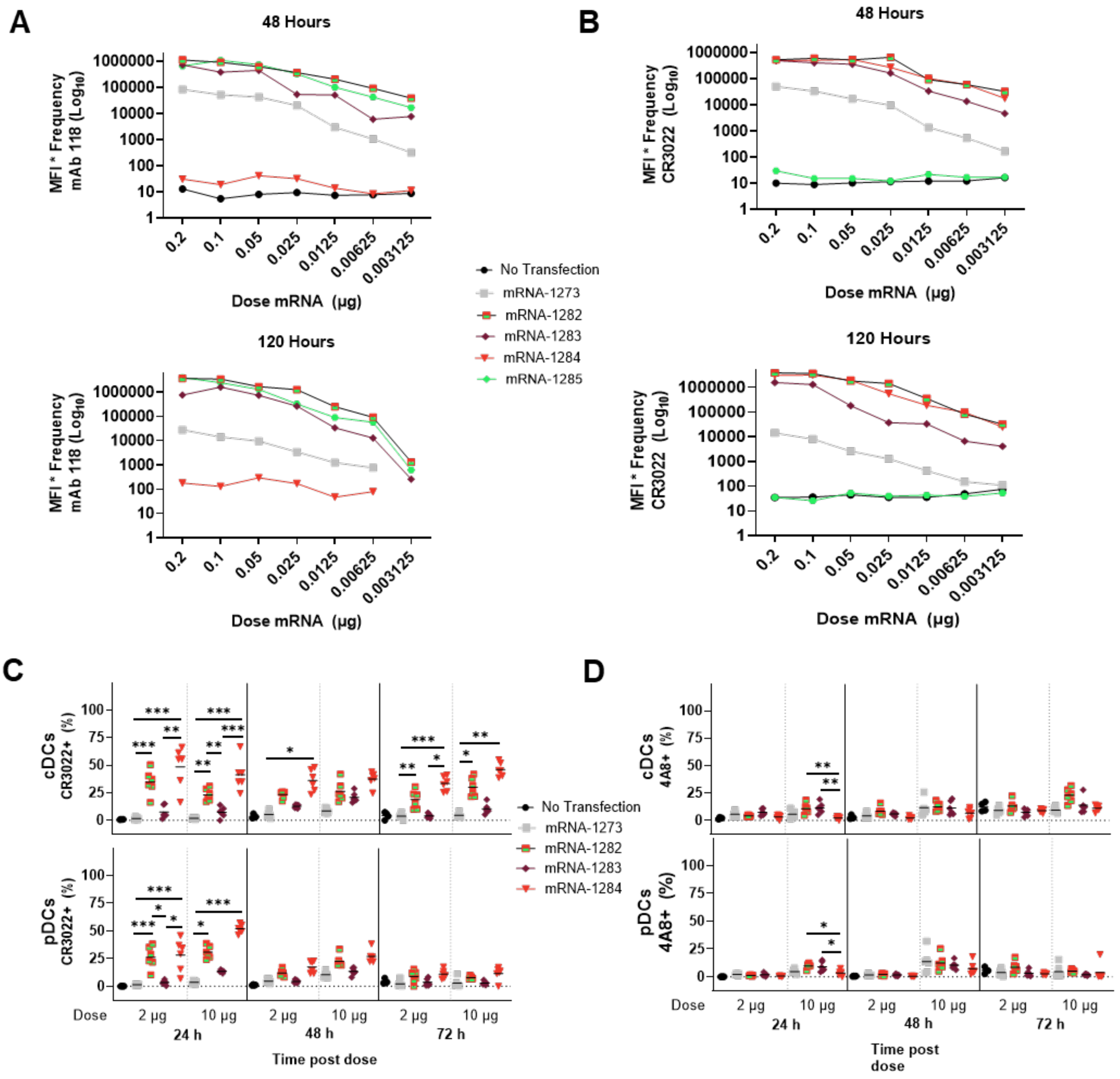

**Figure S2. Immunogenicity and T-cell cytokine responses in mice administered mRNA-1273 and domain-based vaccines**. S-2P specific (left; Panel A only), RBD specific (middle), and NTD-specific (right) IgG titers at Day 21 (**A**) and Day 36 (**B**) for either 2-dose or 1-dose regimens of single mRNA constructs administered at various concentrations. PBS, N=10; 1-µg doses (2 doses), N=12; 0.1-µg doses, N=12; 0.01-µg doses, N=8; 0.001-µg doses N=8; 1-µg dose (1 dose), N=8. (**C**) CD4+ T cells expressing IFN-γ, TNFα, IL-2, IL-4, and IL-5 in response to the S1, S2, and RBD peptide pools. (**D**) CD8+ T cells expressing IFN-γ, TNF-α, and IL-2 in response to the S1, S2, and RBD peptide pools. PBS, N=5; for 1-µg doses: mRNA-1273, N=5; mRNA-1284, N=7; mRNA-1285, N=6; mRNA-1283, N=6; for 0.1 µg doses: mRNA-1273, N=7; mRNA-1284, N=7; mRNA-1285, N=6; mRNA-1283, N=7.

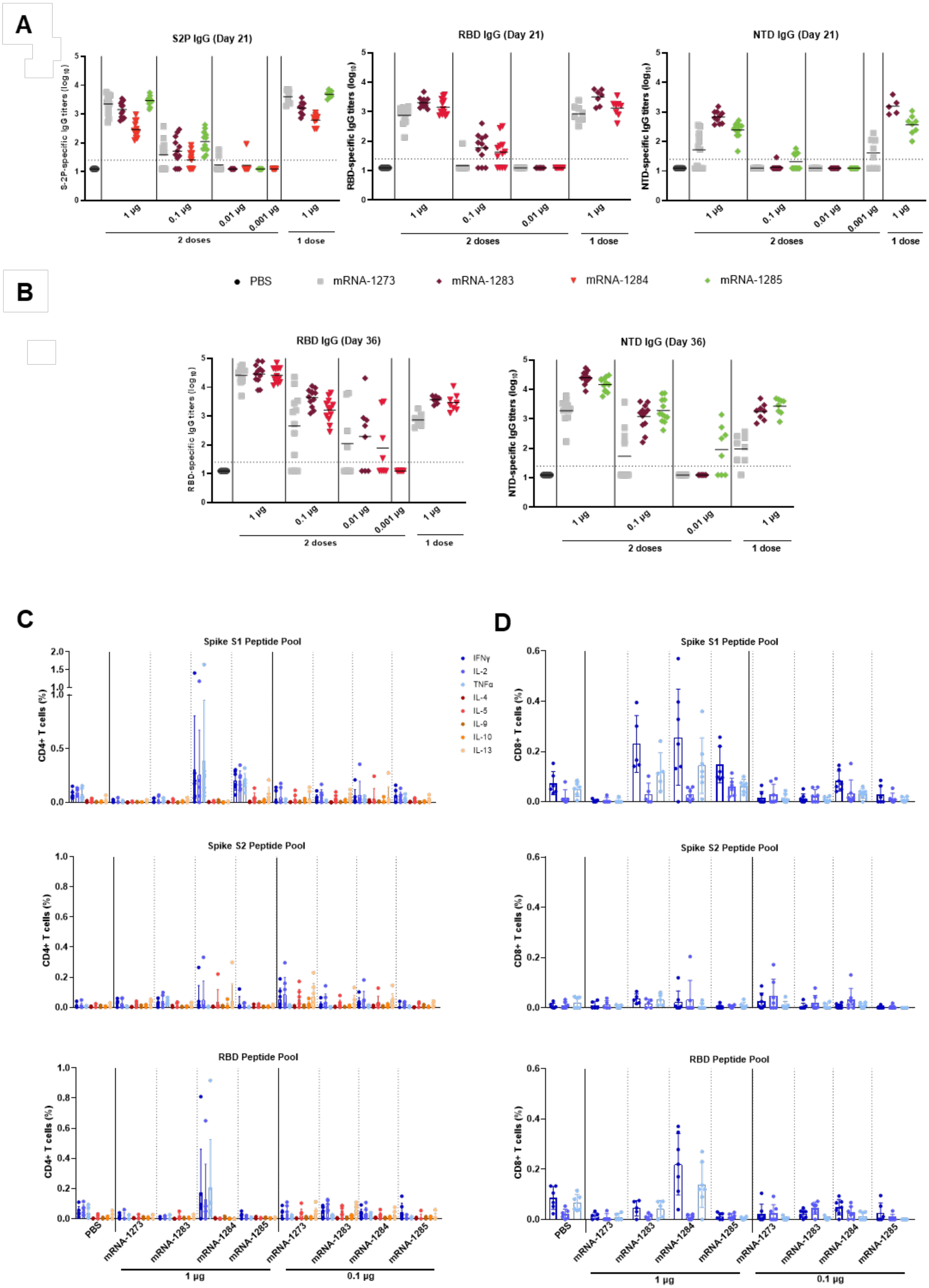
A
S2P IgG (Day 21)
S-2P-specific IgG titers (log10)
RBD IgG (Day 21)
RBD-specific IgG titers (log10)
NTD IgG (Day 21)
NTD-specific IgG titers (log10)
1 µg
0.1 µg
0.01 µg
0.001 µg
2 doses
1 dose
PBS
mRNA-1273
mRNA-1283
mRNA-1284
mRNA-1285
B
RBD IgG (Day 36)
NTD IgG (Day 36)
C
Spike S1 Peptide Pool
Spike S2 Peptide Pool
RBD Peptide Pool
CD4+ T cells (%)
IFNγ
IL-2
TNFα
IL-4
IL-5
IL-9
IL-10
IL-13
D
CD8+ T cells (%)

**Figure S3. Serum neutralization of mice vaccinated with a 2-dose primary series of mRNA-1283 against D614G and BA.1.** (**A-D**) Percentage relative infection of D614G and BA.1 for serum dilutions of mice administered 5 µg or 0.1 µg of the control or (**E-H**) mRNA-1283

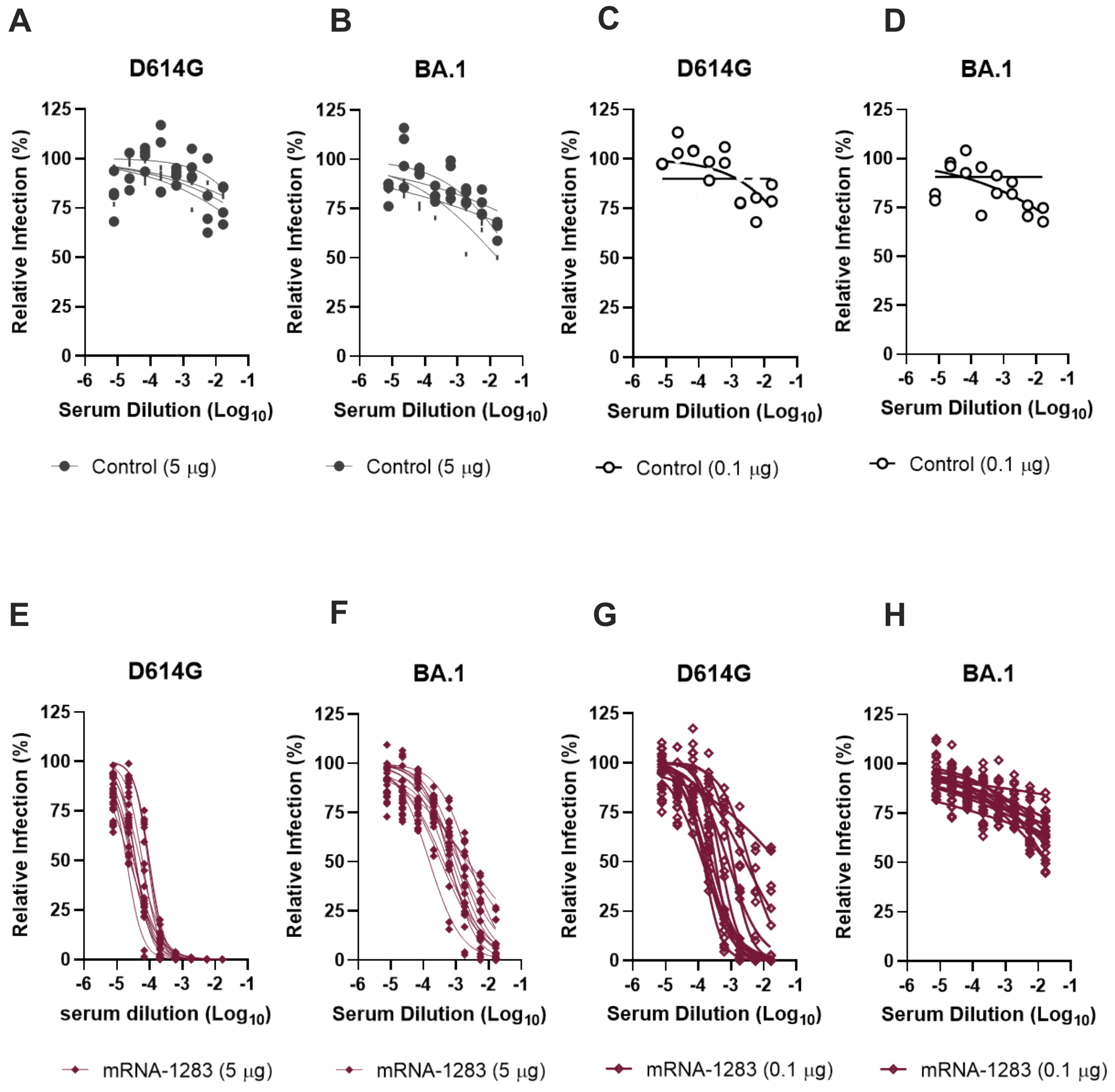

**Figure S4. Nasal wash viral N copies in mRNA-1273– or mRNA-1283–vaccinated mice (0.1- and 5-µg doses) challenged with D614G and BA.1, N=8-10.** Previously published data from control mRNA and mRNA-1273 vaccination were used for reference in this figure, as they were performed concurrently with mRNA-1283 evaluation (23). Multiple comparisons were performed using the Kruskal-Wallis test following calculation of adjusted *P* value via the Dunn's multiple comparison test. ns, non-significant; *, $P<0.05$, **, $P<0.01$, and ***, $P<0.001$.

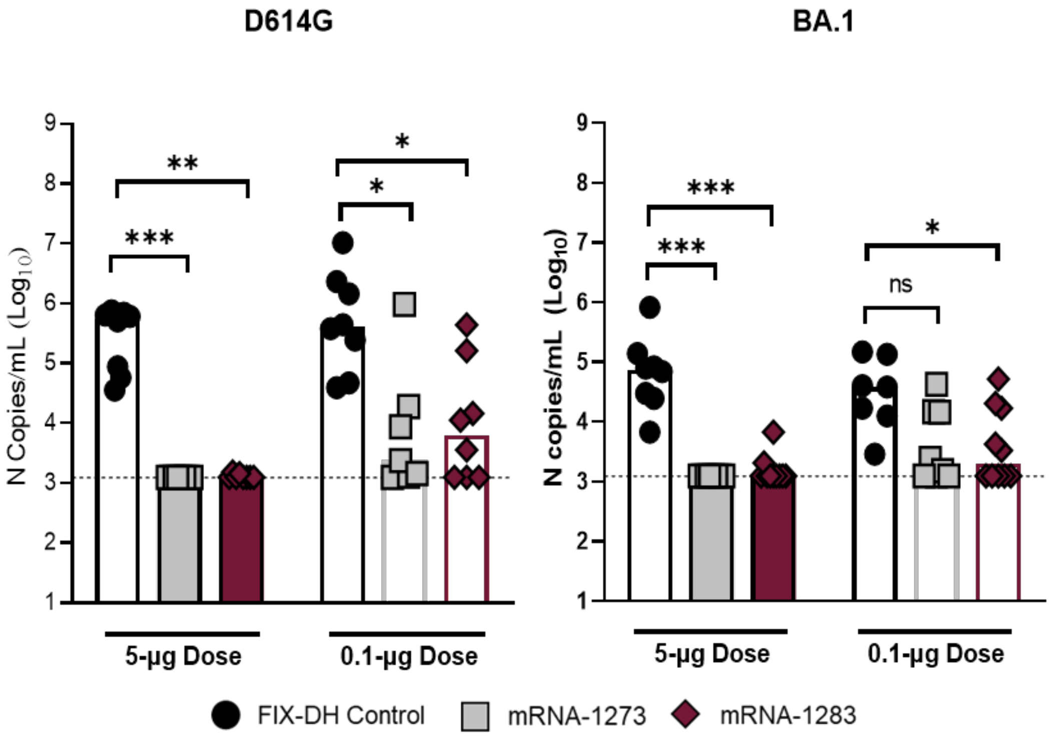
